# Neurobehavioural signatures in race car driving

**DOI:** 10.1101/860056

**Authors:** Ines Rito Lima, Shlomi Haar, Lucas Di Grassi, A. Aldo Faisal

## Abstract

Recent technological developments in mobile brain and body imaging are enabling new frontiers of real-world neuroscience. Simultaneous recordings of body movement and brain activity from highly skillful individuals as they demonstrate their exceptional skills in real-world settings, can shed new light on neurobehavioural structure of human expertise. Driving is a real-world skill which many of us acquire on different levels of expertise. Here we ran a case-study on a subject with the highest level of driving expertise - a Formula E Champion. We studied the expert driver’s neural and motor patterns while he drove a sports car in the “Top Gear” race track under extreme conditions (high speed, low visibility, low temperature, wet track). His brain activity, eye movements and hand/foot movements were recorded. Brain activity in the delta, alpha, and beta frequency bands showed causal relation to hand movements. We demonstrate, here in summary, that even in extreme situations (race track driving) a method for conducting human ethomic (Ethology + Omics) data that encompasses information on the sensory inputs and motor outputs outputs of the brain as well as brain state to characterise complex human skills.

## Introduction

One of the hallmarks of being human is our unique ability to develop skills and expertise. While all animals develop skills like walking, running, fruit peaking or hunting, we as humans can develop much wider and more diverse set of skills. With practice, most of us can learn playing a musical instrument, sports, arts, and crafts. Yet, only some of us can take it to a level of expertise. Unlike the popular view that it is all about practice ^1^, there are accumulating evidence that practice is not enough ^2^. This is probably why we are so fascinated by individual musicians, artists, athletes, and craftsmen who take these expertise to levels that most of us can only imagine. Novel technology for mobile brain and body imaging is now enabling to study neurobehavior in real-world settings ^3^. When carried out in natural environments, these measures can enable a meaningful understanding of human behaviour while performing real-life tasks. Studying of the relation and inter-dependencies between brain activity and body movements of experts, while they perform their expertise in real-world settings, can enable unpacking the enigma of expertise. In recent years there is accumulating body of literature studying the neural signals associated with expertise, particularly in sports ^4^. EEG studies link expert performance to changes in EEG alpha and beta rhythms, however, most of these studies are using lab-based tasks, and therefore their findings have had little impact for sports professionals ^4^. For example, Go/No-Go task was used to study baseball expertise ^5^. While their findings showed that experts perform better and have stronger EEG inhibition responses, which can tell us something about their skill, it is far from the real expertise. Other studies addressed expertise in the real task in a trial-by-trial design. For example, expert rifle shooters exhibited longer quiet eye period before shooting and showed an increased asymmetry in alpha and beta power (increase in left-hemisphere and decrease in right-hemisphere) during the preparatory period ^6^. Also, expert golf players show a greater reduction in EEG theta, high-alpha, and beta power during action preparation ^7^. Measuring and interpreting the neurobehavior of expertise in continues performance of real-world task is the next challenge.

Here we present a case study in real-world neuroscience of expertise, measuring the brain activity and body movement of an expert race car driver, Formula E Champion, while driving under extreme conditions (high speed, low visibility and road slipperiness). Driving is a skill that most of us acquire and use on daily life. Previous literature on driving mostly addressed it as such and thus focused on evaluating brain signals, eye gaze, or body movements, in order to measure the state of attention ^8, 9^, fatigue ^10–12^ and drowsiness ^13, 14^ of the driver, which are major causes of road accidents. Here, we use driving is an exciting test-bed to study real-world expertise. We address comprehensively the driver’s natural perceptual input (vision), motor output (eyes and limbs movements) and brain activity, in attempted to assess the neuromotor behaviour responses under challenging driving conditions which are hallmarks of expertise in race driving.

The objective of this study was to characterise the neuromotor behaviour of expertise in the wild, with an experiment performed in real life conditions, outside a laboratory or simulator, fostering the assessment of a skilled real world task. Better understanding of expert driver’s neural and motor interdependencies while facing driving challenges, can also potentially foster the development of technologies to prevent critical conditions and improve driving safety, as well as safety procedures for autonomous and semi-autonomous cars.

## Methods

### Experimental Setup

The experiment took place on the Dunsfold Aerodrome (Surrey, UK), known as the Top Gear race track. The driver (co-author) was Formula E champion, Lucas Di Grassi (Audi Sport ABT Schaeffler team), with over 15 years of professional racing experience, which include kart, Formula 3, Formula One and Formula E racing. The test drive was prescheduled for video production purposes by the racing team, who race in these conditions frequently. It enabled us this unique scientific observation of motor expertise in the wild. Although unlikely needed, emergency response units were present. A promo video of the film by Averner Films^15^ is accessible here: vimeo.com/248167533.

The driver was equipped with: a 32-channels wireless EEG system (LiveAmp, Brain Products GmbH, Germany) with dry electrodes (actiCAP Xpress Twist, Brain Products GmbH, Germany); binocular eye-tracking glasses (SMI ETG 2W A, SensoMotoric Instruments, Germany); and four inertial measurement units (IMUs) on his hands and feet (MTw Awinda, Xsens Technologies BV, The Netherlands); shown in Figure 1A. The car was equipped with a GPS system and a camera recording the inside of the car. The car driver assistance systems were turned off. The full architecture of the experimental setup is presented in Figure 1C.

**Figure 1.**
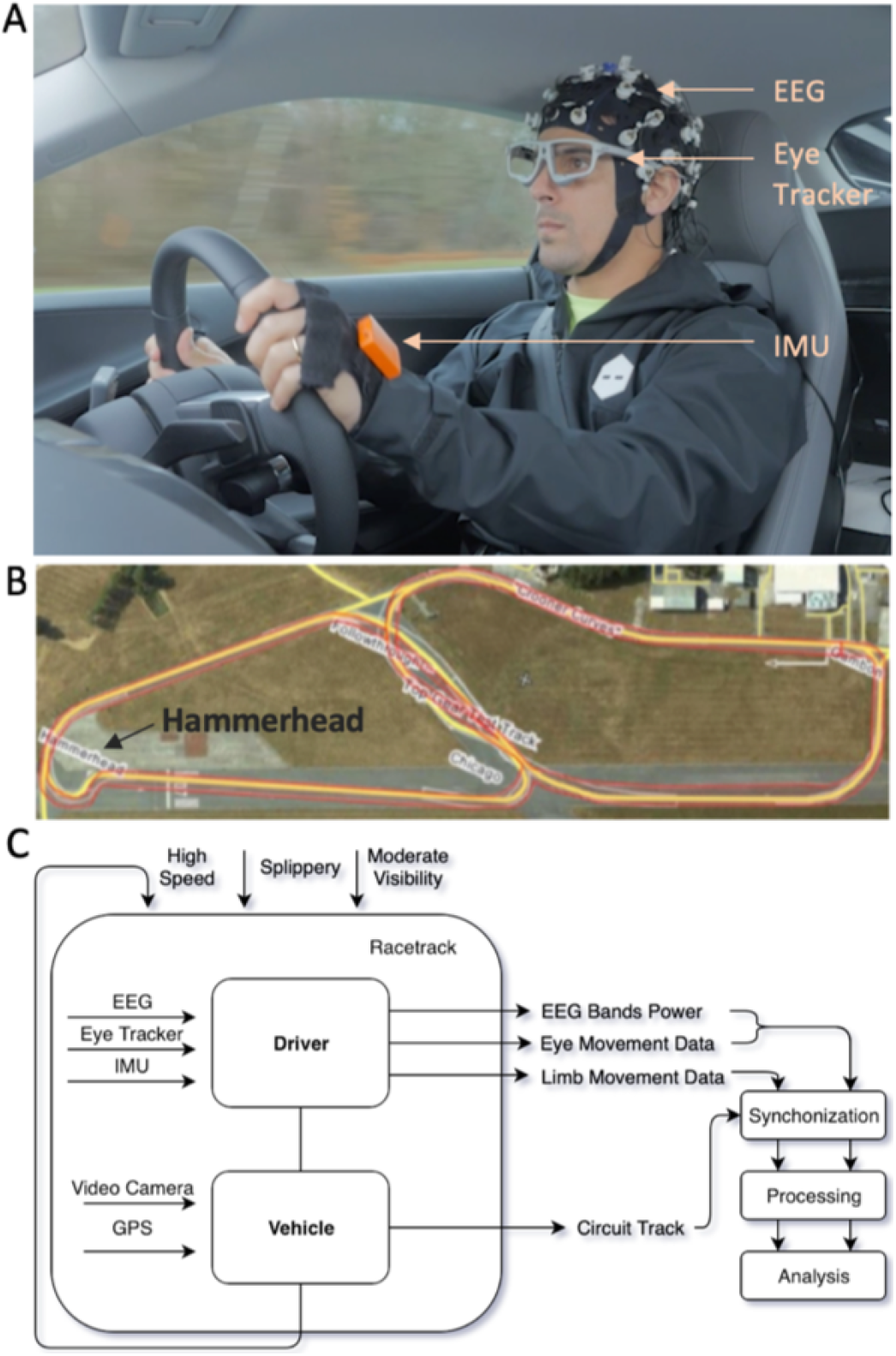
Experimental setup involving expert driver (LDG) : A) involved equipment (EEG, eye tracking and body motion tracking systems) and their placement on driver; B) Top Gear racetrack and the highlighted race critical curve, the *Hammerhead*; C) architecture of our data collection system with driver/car/road parameters inputs and data flows.

The Top Gear race track has specific curve types that subjugate drivers’ to different driving challenges. In particular, in the south-west of the track (left side of of the image in Figure 1B) is the *Hammerhead* curve. As the name suggest, it is a hammerhead-shaped tricky curve, designed to test cars and the driver skills for it is technically challenging^16^. This is considered the critical curve of the track, being used in this work to asses the driver’s performance in challenging scenarios. This extreme curve was selected to show the clear distinguished driver neuromotor behaviour.

The car was driven with a mean driving speed of 120 Km/h and a top speed of 178 Km/h on a track 2.82 Km long with 12 curves. The experiment consisted of 6 sets of 2 to 3 laps each, stopping after every set for system check up and recalibration.

### Data Processing

All data pre-processing used custom software written in MatLab (R2017a, The MathWorks, Inc., MA, USA). Timestamp were synchronized across all data streams using cross-correlation between the accelerations recorded by all system, then data streams were segmented to the laps and laps with missing data in either stream were removed.

#### Car movement data

The car GPS was obtained by the placement of an iPhone inside the car. This system’s acceleration and rotation recordings were resampled to a stable 100Hz sampling rate, for timestamp synchronization and for cleaning the body data. The data was filtered resorting to a zero phase-lag fourth-order Butterworth filter with 10Hz cut off frequency ^17, 18^.

#### Body movement data

Body data was collected by 4 Xsens MTw Awinda IMUs placed on hands and feet. These sensors recorded the rotation and acceleration. The data collected has a stable sampling rate of 100Hz. The motion captured by the motion caption system, is the combination of the car movement and the drivers limbs movements inside the car. A linear regression was implemented in order to clean the car movement (from the GPS recordings) from the motion tracking. The regression’s residuals capture the limbs movement that does not fit the car movement, for rotation and acceleration data separately. The data was filtered resorting to a zero phase-lag fourth-order Butterworth filter with 10Hz cut off frequency ^17, 18^.

#### Gaze data

The eye gaze was acquired with binocular SMI ETG 2W A eye-tracking glasses which use infrared light and a camera to track the position of the pupil. Using the pupil centre and the corneal reflection (CR) creates a vector in the image that can be mapped to coordinates^19, 20^. The glasses egocentric camera captures the scene view and enabling the reconstruction of the frame-by-frame gaze point in the real-world scene. The gaze data was collected at 120Hz and included multiple standardised eye measurements, such as pupil size, eye position, point of regard, and gaze vector. Our analysis focused on the point of regard, obtained from the RMS between the point of regard binocular measured in X and Y axis; the gaze vector, computed for the right and left eyes, separately, through the RMS of XYZ; and the change in gaze vector, calculated using two consecutive measurements. These measures were resampled to 100Hz and linear interpolation was applied to account for some missing data points in the gaze recording.

#### EEG data

The brain activity was recorded using a 32 channels EEG cap with dry electrodes (displayed accordingly to the 10-20 system), sampled at 250Hz stable sampling rate. EEG data was analysed with EEGLAB toolbox (https://sccn.ucsd.edu/eeglab;^21^). The processing steps included i) high pass filtering with cutoff frequency of 1Hz^22–24^; ii) line noise removal in the selected frequencies of 60Hz and 120Hz^22^; iii) bad channels removal for those with less than 60% correlation to its own reconstruction based on its neighbour channels^22^; iv) re-referencing the EEG dataset to the average (of all channels) to minimise the impact of a channel with bad contact in the variance of the entire dataset^22^; v) Independent Component Analysis (ICA) in order to separate signal sources^25–27^; vi) Artifacts removal using *runica, infomax* ICA algorithm from EEGLAB, for identification and removal of head and eye movements and blinking artifacts. EEG data was transformed in the time-frequency domain to power in decibels (dB) in 100Hz (to match the other data streams), in *delta* (*δ*) 0.5 to 4Hz; *theta* (*θ*) 4 to 8Hz; *alpha* (*α*) 8 to 12Hz; and *beta* (*β*) 12 to 30Hz. frequencies bands. The transformation was applied separately to the individual IC located over the left motor cortex and to the mean brain activity, averaged across the cleaned ICs.

## Results

Initial analysis was aimed to understand the interdependencies between the different neurobehavioural data streams and their level of complexity. The analysis then focus on specific driving events, such as response to challenging conditions (skidding, curves and straights), in order to assess if there is a distinguishable behaviour upon those moments. Lastly we addressed the causality across different data streams.

### Data characteristics

The distributions of the car speed, the right hand rotation, and the right hand accelerations were assessed in the entire track, the straight segments before and after the hammerhead curve, and the hammerhead curve itself (Figure 2). The average speed throughout the experiment was 120Km/h. As expected, in the curves speed was relatively low (between 54 and 82 Km/h), while in straight segments was much higher with a wider distribution, as the car decelerated towards a curve and accelerated after the curve. The number of frames considered hammerhead critical curve was 11.3% of the total recording and straights corresponds to 25.5%.

**Figure 2.**
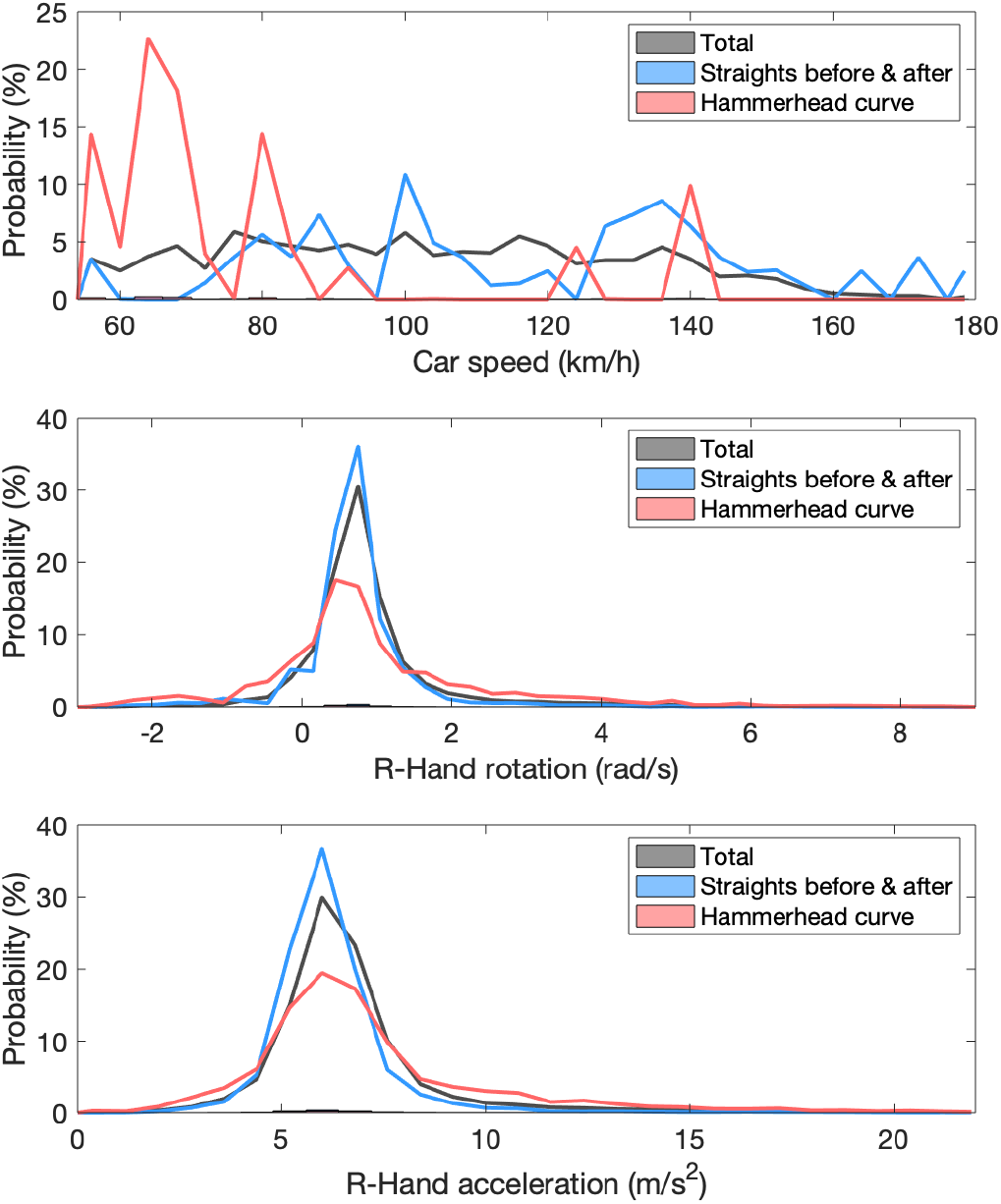
Histogram evolution analysis detail the extreme driving scenario of this experiment regarding: (top) car speed with and average of 120Km/h, critical curve speed average of 78Km/h and straight speed average of 130Km/h, all above conventional driving speed limits; (middle) right hand gyroscope with average value of 0.9 rad/s whereas for intense driving style (critical curve and skidding moments) the average values are of 1 and 3 rad/s, superior to normal movement expected from literature; and (bottom) right hand accelerometer data showing absolute acceleration with similar results distribution as gyroscope data. Data regarding straights corresponds to 25.5% of the entire data set and the Hammerhead curve to 11.3%.

Since the hand movements were highly correlated here we show only the right hand. Both gyroscope and accelerometer distributions present similarly tendencies, with a narrow values distribution during the straight segments and a slightly spreader one in curves. The result considering the total data set lies in-between.The gyroscope values for the abrupt responses (skidding) have a mean of 3rad/s, considerably superior to the 1rad/s found in literature for normal forearm movement^28^, which is expected considering the intense car handling. Data considered as abrupt responses corresponds to 6% of the dataset.

Eye gaze data showed a strong tendency of tracking the tangent point of the curve, as illustrated in Figure 3 (top), and reported for non-expert drivers ^29^. In the top figure can be seen the eye gaze search for the tangent of the curve (internal and external), marked in white on the road, where it remains throughout the entire curve. During straight segments, the eye gaze focus straight ahead, with a stable distance in the horizon, with minor saccadic deviations, as illustrated in Figure 3 (bottom). The heat map was built using data recorded during the critical curve (top) or during the straights before and after that curve (bottom). The gaze point position from the egocentric view was annotated at a 10 frames cadence in a overlapping position matrix. The heat map was built resorting to the percentual annotations occurrence in this matrix.

**Figure 3.**
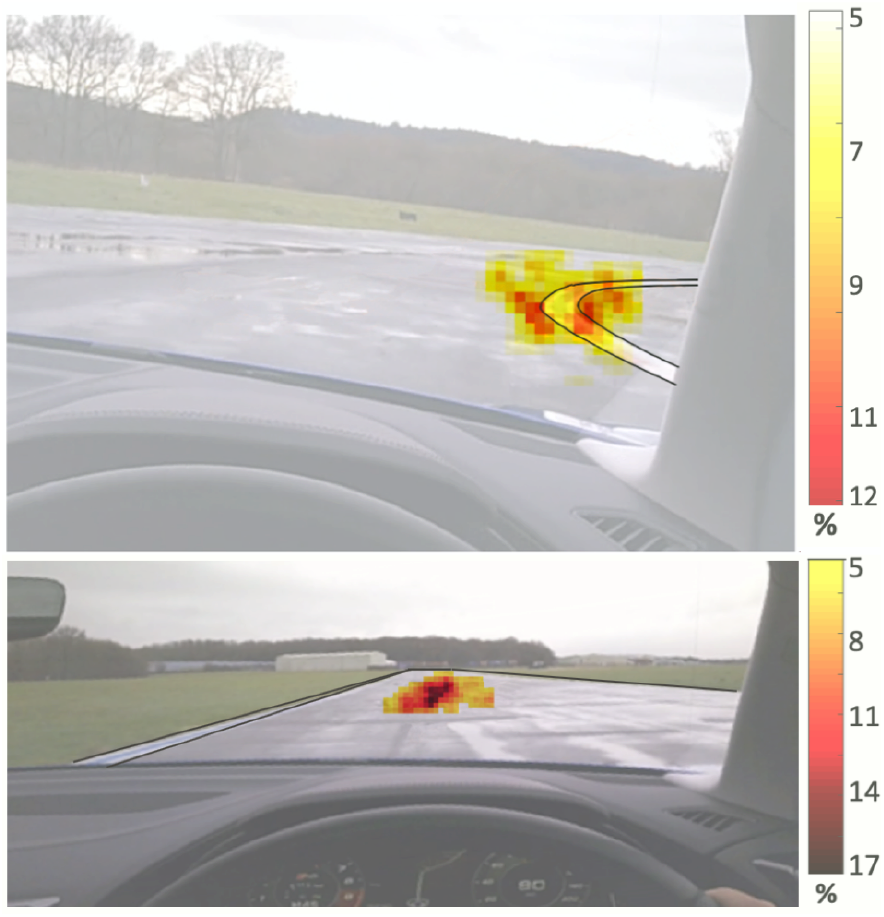
Heat maps of eye gaze using data recorded during: the *Hammerhead* critical curve (top), being visible the tendency of tracking the tangent of the curve; and the straight segments before and after the curve (bottom), where the gaze focus on the horizon ahead is visible.

### Global dataset assessment

For an overall understanding of the interdependencies between the neurobehavioral data streams, a correlation matrix was computed (Figure 4A). The variables considered were: the RMS of the acceleration and the rotation over the four limbs, the car, and the head; EEG band power for the average brain activity, and for the right hand IC; and eye movement data including the point of regard, the gaze vectors, and the change in gaze vector for both eyes. The right and the left hands are strongly correlated, as expected since the driver had both hands on the steering wheel. There are other correlation within domains (e.g. neighboring band powers are correlated), and also weaker, but statistically significant, correlations between domains. Most importantly, between the EEG band powers and the hand movements. We found a positive correlation for the hands acceleration and rotation with the alpah and beta power in the right hand IC, and a negative correlation with the delta power (P<0.001). Cross-correlation shows that while the delta band is synced with the movement the neural signal in the alpha and beta band precedes the movement by 100ms.

**Figure 4.**
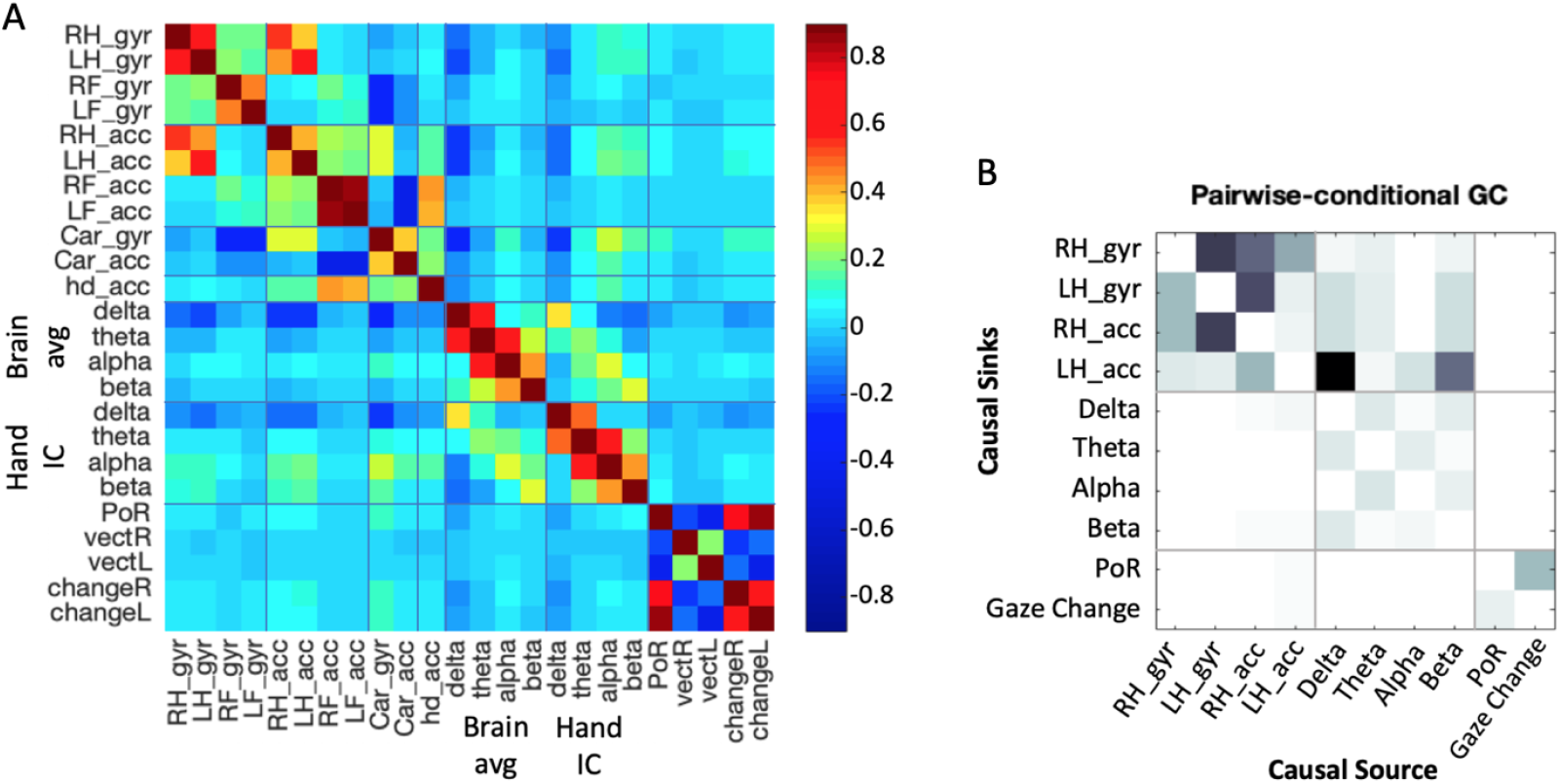
Global assessments: A) Instantaneous correlation matrix between all data groups: gyroscope rotation (gyr) and acceleration (acc) of the right and left hands (RH,LH), the right and left feet (RF,LF), the car (Car), and the head (hd); the band power in the delta, thata, alpha, and beta bands across the brain (Brain avg) and in the left motor cortex (Hand IC); eye gaze point of regard (PoR), the gaze vectors (vect) and the change in gaze vector (change) for the right (R) and the left (L) eye; B) Granger causality pairwise results considering the data groups separated by grey lines. Darker color is associated with stronger causality, for which brain data is more cause of hand movements than the other way around, whereas eye data does not cause any of the variables and is barely caused by *alpha* and *beta* bands power. A significance of p=0.05 and Bonferroni correction supported these conclusions.

#### Granger causality

To address the sequence of causality between variables we used Granger causality using EEGLAB toolbox MVGC multivariate Granger causality (mvgc_v1.0) ^30, 31^. Granger causality analysis was conducted on the hands acceleration and gyroscope rotation data, EEG bands power from the IC of the right hand, the gaze point of regard, and the RMS across right and left eye gaze change vector. The results are presented in Figure 4B as a pairwise matrix of causality between causal sources to causal sinks (darker indicates stronger causality) for statistically significant causality links (p>0.05 after Bonferroni correction). We found one directional causality where the EEG bands power are causing hand movements, specially *delta* and *beta*. Also, the eye data is not classified as cause or effect of hand movements or low frequency EEG power bands.

### Specific driving related events

Here is considered a comparison between the neurobehavioral signature of the challenging Hammerhead curve and the straight segments leading to it and out of it. Based on the GPS data and the egocentric videos we annotated the segments of the critical *Hammerhead* curve in the track and the straight paths leading to it and out of it. Correlation matrix from hands IMUs and EEG data streams for these segments shows differences in the correlation structure between the two types of segments (Figure 5A). While in the straight segments there was no correlation between the hands’ rotation and acceleration to the EEG power, in the curve there was a negative correlation between these brain and body measurements. Statistical analysis of the differences between the segments shows a decrease in *delta* power and an increase in *alpha* and *beta* during the curve (Figure 5B). The data from before and after the curve also showed differences where *delta* power is lower before and higher after the curve, while *alpha* power shows the opposite trend. Eye gaze vector also changed more during the curves.

**Figure 5.**
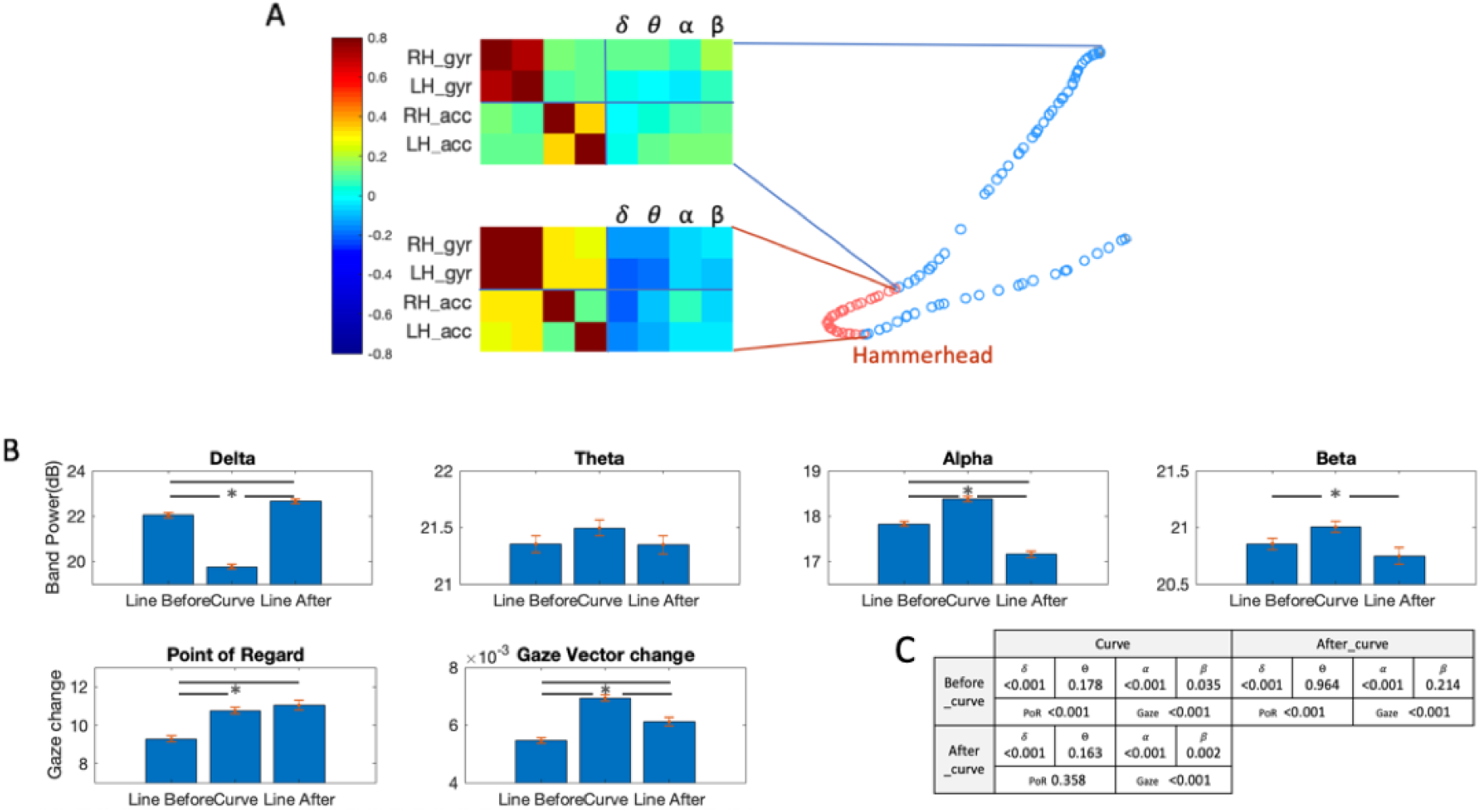
A) Identification of the straights (blue) and curve (red) periods, with a fragment of the correlation matrix obtained for each period; B) statistical analysis on the mean and SEM for curves and straights on bands power and eye data. Despite of lower correlation values, the brain shows structured correlation to the hand when performing the curve, with *delta* and *theta* anticorrelation and *alpha* and *beta* positive correlations, instead of a generalized correlation to the body. Statistical significance between before, during and after the critical curve was found for *delta*,*alpha*,*beta*, point of regard and change in gaze vector. C) shows the p-values obtained with t-test

## Discussion

This work is a first attempt to assess neurobehavioural signatures of real-world motor expertise in-the-wild. We demonstrate the feasibility of simulatenously measuring brain activity, eye and body movement behaviours in-the-wild under extreme conditions (race track driving) and assessing neurobehavioural interdependencies and infer causal relations. Our results show changes in the EEG power and the gaze characteristic during sharp curves, where our driver’s and car control expertise are most likely to make a difference. Interestingly, the EEG power changes are inline with previous results on general creative solution finding and interventions ^32^.

Comparing the driver’s neurobehaviour betweeen the sharp curves and the straight segments before and after, enabled us to assess world-championship-level skilled responses while demonstration their expertise. Our results suggest clear differences in the EEG power, point of regard, and gaze change vector, between the different segments: before-during-after sharp curves. During the curves, the driver showed a power increase in the alpha and beta bands and a decrease in the delta band. The increase in alpha is potentially a signature of the increased skillful creativity demand in these segments ^32, 33^. The alpha and beta power increase we observe are also inline with previous work showing increase in left-hemisphere alpha and beta power of expert rifle shooters during the preparatory pre-shot period ^6^.

Neurobehavioural data collection in-the-wild is subject to more noise sources and interference than standard data collection in-the-lab. This concern is specifically worrying for the EEG signal, which is always contaminated by noise, and any EEG recording during movement it is subject to movement and muscle artifacts. Thus, we find our cross-correlation and Granger causality results very encouraging, as those suggest the the EEG activity precedes the movement and predicted it and not the other way around. Therefore, the EEG results cannot be rejected as noise artifacts.

The driver’s gaze during curves followed the tangent point of the curve, as suggested in the classic paper by Land and Lee ^29^. During the straight segments his gaze was perfectly focused on the center of the road which led to the more stability in straight segments relative to curves, though during both segments type the driver’s gaze is extremely stable, as illustrated in Figure 3.

Being able to collect real-world data which capture a significant portion of the sensory input to the brain (visual scene and locus of attention), the motor output of the brain (hand, head and arm movements during driving) as well as the state of the brain (EEG signals), is a further realisation of our human ethomics approach. This is not only insightful for understanding the brain and its behaviour, but also for devising artifical intelligences. In recent work^34^ we demonstrated how human drivers in a virtual reality driving simulator generated gaze behaviour that we used to train a deep convolutional neural network to predict where a human driver was looking moment-by-moment. This human-like visual attention model enabled us to mask “irrelevant” visual information (where the human driver was not looking) and train a self-driving car algorithm to drive successfully. Crucially, our human-attention based AI system learned to drive much faster and with a higher end-of-learning performance than AI systems that had no knowledge of human visual attention. The work we present here takes this *en passant* human-annotation of skilled beahviour to the next level, by collecting real-world data of rich input and output characteristics of the brain. Similarly, on the side of controlling system we showed for example how using ethomic data obtained from natural tasks (movement data^35^, electrophysiological data^36^, decision making data^37^) can be harnessed to boost AI system performance. The neurobehavioural approach demonstrated here suggests how we can in future close-the-loop fully between person and vehicle.

In summary, we demonstrated the feasibility to study the neurobehavioral signatures of real-world expertise in-the-wild. We showed evidence of specific brain activity and gaze patterns related to expert’s extreme driving, in particular during driving events in which the expertise of the driver makes a key difference.

## Acknowledgements

First we would like to acknowledge Alex Verner and Joel Verner (Averner Films) who’s video production idea and work enabled this research. We thank them and the race team for inviting us to participate in their production and collect the data for this study. We also thank Pavel Orlov for his help with the gaze data. The study was enabled by financial support to a Royal Society-Kohn International Fellowship (NF170650; SH and AAF) and by eNHANCE (http://www.enhance-motion.eu) under the European Union’s Horizon 2020 research and innovation programme grant agreement No. 644000 (SH and AAF).

## Author contributions statement

AAF conceived and designed the scientific study; LDG preformed the experiment; IRL and AAF acquired the data; IRL, AAF and SH analyzed the data; IRL, SH and AAF interpreted the data; IRL and SH drafted the paper; IRL, SH, LDG and AAF revised the paper.

## Additional information

The experimental data was collected and analysed here was as part of a promotional video production for Roborace - a racing competition for autonomously driving, electrically powered vehicles. LDG is the CEO of Roborace. Averner Films was comissioned by Roborace to produce the video. LDG and AAF appeared in the video. IRL and SH declare no competing financial interests. Within the domain of this paper AAF has consulted for Airbus, Averner Films and Celestial Group. AAFs lab has received within the domain of this paper research funding and donations from Microsoft and NVIDIA. This scientific publication was not commissioned nor expected as part of the film production.

